# Unsupervised extraction of stable expression signatures from public compendia with eADAGE

**DOI:** 10.1101/078659

**Authors:** Jie Tan, Georgia Doing, Kimberley A. Lewis, Courtney E. Price, Kathleen M. Chen, Kyle C. Cady, Barret Perchuk, Michael T. Laub, Deborah A. Hogan, Casey S. Greene

## Abstract

Cross experiment comparisons in public data compendia are challenged by unmatched conditions and technical noise. The ADAGE method, which performs unsupervised integration with neural networks, can effectively identify biological patterns, but because ADAGE models, like many neural networks, are over-parameterized, different ADAGE models perform equally well. To enhance model robustness and better build signatures consistent with biological pathways, we developed an ensemble ADAGE (eADAGE) that integrated stable signatures across models. We applied eADAGE to a *Pseudomonas aeruginosa* compendium containing experiments performed in 78 media. eADAGE revealed a phosphate starvation response controlled by PhoB. While we expected PhoB activity in limiting phosphate conditions, our analyses found PhoB activity in other media with moderate phosphate and predicted that a second stimulus provided by the sensor kinase, KinB, is required for PhoB activation in this setting. We validated this relationship using both targeted and unbiased genetic approaches. eADAGE, which captures stable biological patterns, enables cross-experiment comparisons that can highlight measured but undiscovered relationships.

## Introduction

Available gene expression data are outstripping our knowledge about the organisms that we’re measuring. Ideally each organism’s data reveals the principles underlying gene regulation and consequent pathway activity changes in every condition in which gene expression is measured. Extracting this information requires new algorithms, but many commonly used algorithms are supervised. These algorithms require curated pathway knowledge to work effectively, and in many species such resources are biased in various ways (Gillis and Pavlidis, 2013; Greene and Troyanskaya, 2012; Schnoes et al., 2013). Annotation transfer can help, but such function assignments remain challenging for many biological processes (Jiang et al., 2016). An unsupervised method that doesn’t rely on annotation transfer would bypass the challenges of both annotation transfer and biased knowledge.

Along with our wealth of data, abundant computational resources can now power deep unsupervised applications of neural networks, which are powerful methods for unsupervised feature learning (Bengio et al., 2013). In a neural network, input variables are provided to one or more layers of “neurons”. Each neuron (also called node) has an activation function that determines whether or not it turns on given some input. The entire network is trained, which consists of adjusting the edge weights that each node provides to each other, by grading the quality of the output for some task. Denoising autoencoders (DAs), a type of unsupervised neural networks, are trained to remove noise that is intentionally added to the input data (Vincent et al., 2008). Masking noise, in which a fraction of the inputs are set to zero, is commonly used (Vincent et al., 2010) and successful denoising autoencoders must learn dependency structure between the input variables. Adding artificial noise helps a DA to learn features that are robust to partial corruption of input data. This approach has properties that make it particularly suitable for gene expression data (Tan et al., 2015). First, the sigmoid activation function produces features that tend to be on or off, which helps to describe biological processes, e.g. transcription factor activation, with threshold effects. Second, the algorithm is robust to noise. We previously observed that a one-layer DA-based method, ADAGE (analysis using denoising autoencoders of gene expression), was more robust than linear approaches such as ICA or PCA in the context of public data, which employ heterogeneous experimental designs, lack shared controls, and provide limited metadata (Tan et al., 2016b).

Neural networks have many edge weights that must be fit during training. Given some gene expression dataset, there are many different DAs that could reconstruct the data equally well. In a technical sense we would say that the objective functions of neural networks are typically non-convex and trained through stochastic gradient descent. When we train multiple models, each represents a local minimum. Yu recently emphasized the importance of patterns that are stable across statistical models in the process of discovery (Yu, 2013). While run-to-run variability obscures some biological features within individual models, stable patterns across neural networks may clearly resolve biological pathways. To directly target stability, we introduce an unsupervised modeling procedure inspired by consensus clustering (Monti et al., 2003). Consensus clustering has become a standard part of clustering applications for biological datasets. Our approach builds an ensemble neural network that captures stable features and improves model robustness.

To apply the neural network approach to compendium-wide analyses, we first sought to create a comprehensive model in which biological pathways were successfully learned from gene expression data. We adapted ADAGE (Tan et al., 2016b) to capture pathways more specifically by increasing the number of nodes (model size) that reflect potential pathways from 50 to 300, a size that our analyses indicate the current public data compendium can support. We then built its ensemble version (eADAGE) and compared it with ADAGE, PCA, and ICA. While it is impossible to specify a *priori* the number of true biological pathways that exhibit gene expression signatures, we observed that eADAGE models produced gene expression signatures that corresponded to more biological pathways. This indicates that this method more effectively identifies biological signatures from noisy public data. While ADAGE models reveal biological features perturbed within an experiment, the more robust eADAGE models also enable analyses that cut across an organism’s gene expression compendium.

To assess the utility of the eADAGE model in making predictions of biological activity, we applied it to the analysis of the *Pseudomonas aerguinosa* gene expression compendium which included 1051 samples grown in 78 distinct medium conditions, 128 distinct strains and isolates, and dozens of different environmental parameters. After grouping samples by medium type, we searched for eADAGE-defined signatures that differed between medium types. This cross-compendium analysis identified five media that elicited a response to low-phosphate mediated by the transcriptional regulator PhoB, and only one of these five media was specifically defined as a condition with low phosphate. While PhoB is known to respond to low phosphate through its interaction with PhoR in low concentrations (Wanner and Chang, 1987), our analyses indicated that PhoB is also active at moderate phosphate concentrations. Specifically, in media with moderate phosphate concentrations, the eADAGE model predicted a previously undiscovered role for KinB in the activation of PhoB, and our molecular analyses of *P. aeruginosa* confirmed this prediction. Analysis of a collection of P. aeruginosa mutants defective in kinases validated the specificity of the KinB-PhoB relationship.

In summary, eADAGE more precisely and robustly captures biological processes and pathways from gene expression data than other unsupervised approaches. The signatures learned by eADAGE support functional gene set analyses without manual pathway annotation. The signatures are robust enough to enable biologists to identify not only differentially active signatures within one experiment, but also cross-compendium patterns that reveal undiscovered regulatory mechanisms captured within existing public data.

## Results

### eADAGE: ensemble modeling improves the model breadth, depth, and robustness

ADAGE is a neural network model. Each gene is connected to each node through a weighted edge (Figure 1A). We define a gene signature learned by an ADAGE model as a set of genes that contribute the highest positive or highest negative weights to a specific node (Figure 1B, see methods for detail). Therefore, one node results in two gene signatures, one on each high weight side. The positive and negative signatures derived from the same node do not necessarily compose inversely regulated processes (Figure S1), so we use them independently.

**Figure 1:**
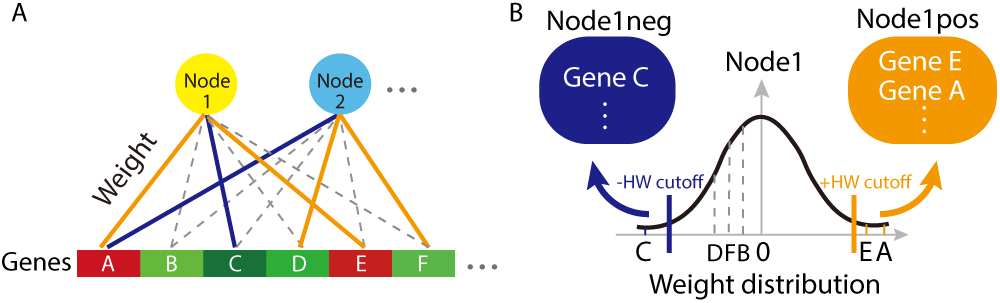
ADAGE model and signature definition. A In an ADAGE model, every gene contributes a weight value to every node. The strength of weight values is reflected by gene-node edge. Orange edges indicate high positive weight. Blue edges indicate high negative weight. Dotted edges show low positive or negative weights. B The distribution of a node’s weight matrix (Node1 as an example) is roughly normally distributed and centered at zero. Genes with weights higher than the positive high-weight (HW) cutoff (GeneE and GeneA) form the gene signature Node1pos. Similarly, genes with weights lower than the negative HW cutoff (GeneC) form the gene signature Node1neg.

ADAGE models of the same size capture different pathways. This occurs because each ADAGE model is initialized with random weights, and the training processes are sensitive to initial conditions. eADAGE, in which we built an ensemble version of individual ADAGE models, took advantage of this variation to enhance model robustness. Each eADAGE model integrated nodes from 100 individual ADAGE models (Figure 2A). To unite nodes, we applied consensus clustering on nodes’ weight vectors because the weight vector captures both the genes that contribute to a node and their magnitude. Our previous ADAGE analyses showed that genes contributing high weights characterized each node’s biological significance, so we designed a weighted Pearson correlation to incorporate gene weights in building eADAGE models (see methods). We compared eADAGE to two baseline methods: individual ADAGE models and corADAGE, which combined nodes with an unweighted Pearson correlation. For direct comparison, the model sizes of ADAGE, eADAGE, and corADAGE were all fixed to 300 nodes, which we found to be appropriate for the current *P. aeruginosa* expression compendium through both data-driven and knowledge-driven heuristics (see supplemental information).

**Figure 2:**
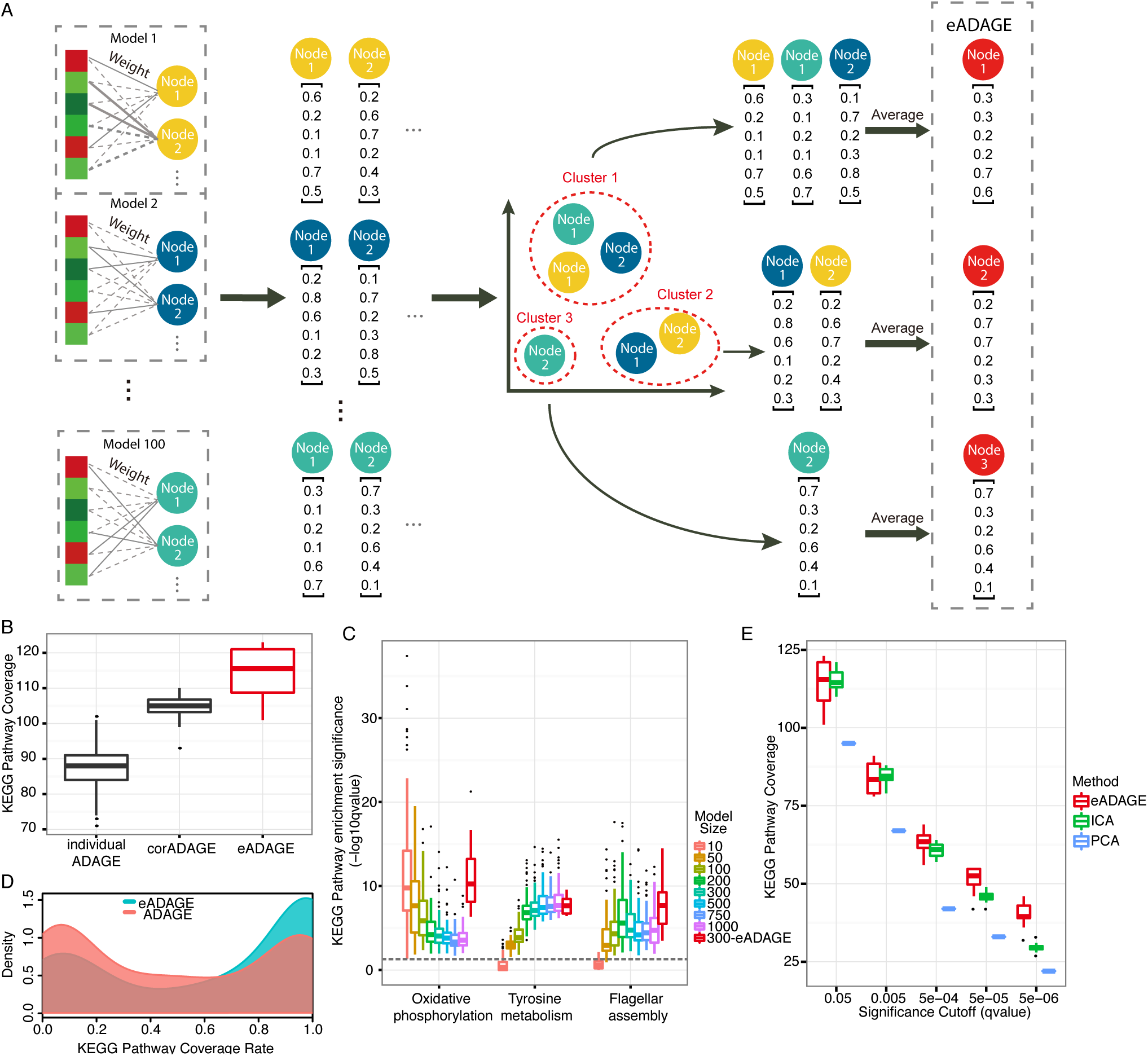
The construction and performance of eADAGE. A eADAGE construction workflow. 100 individual ADAGE models were built using the same input dataset (step 1). Nodes from all models were extracted (step 2) and clustered based on the similarities in their associated weight vectors (step 3). Nodes derived from different models were rearranged by their clustering assignments (step 4). Weight vectors from nodes in the same cluster were averaged and thus becoming the final weight vector of a newly constructed node in an eADAGE model (step5). B KEGG pathway coverage comparison between individual ADAGE and ensemble ADAGE. eADAGE models (n=10) covers significantly more KEGG pathways than both corADAGE (n=10) and ADAGE (n=1000). C The enrichment significance of three example KEGG pathways in different models. The three pathways show different trends as model size increases in individual ADAGE, however, their median significance levels in eADAGE are comparable or better than all individual models with different sizes. The grey dotted line indicates FDR q-value of 0.05 in pathway enrichment. D The distribution of KEGG pathway coverage rate of ADAGE (n=1000) and eADAGE (n=10). eADAGE models have larger proportion on the high coverage side than ADAGE models, indicating pathways were captured more robustly in eADAGE. E Comparison among PCA, ICA, and eADAGE in KEGG pathway coverage at different significance levels. eADAGE outperforms PCA at all significance levels. eADAGE and ICA show similar pathway coverage at the cutoff q-value = 0.05. However, ICA covers less pathways than eADAGE as the significance cutoff becomes more stringent.

While ADAGE models are constructed without the use of any curated information such as KEGG (Kanehisa and Goto, 2000) and GO (Ashburner et al., 2000), we evaluate models by the extent to which they cover the pathways and processes defined in these resources to see how they capture existing biology. For each method, we determined the number of KEGG pathways significantly associated with at least one gene signature in a model, referred to as KEGG coverage. eADAGE models exhibited greater KEGG coverage than those generated by other methods (Figure 2B). Both corADAGE and eADAGE covered significantly more KEGG pathways than ADAGE (t-test p-value of 1.04e-6 between corADAGE (n=10) and ADAGE (n=1000) and t-test p-value of 1.41e-6 between eADAGE (n=10) and ADAGE (n=1000)). Moreover, eADAGE models covered, on average, 10 more pathways than corADAGE (t-test p-value of 1.99e-3, n=10 for both groups). Genes that participate in multiple pathways can influence pathway enrichment analysis, a factor termed pathway crosstalk (Donato et al., 2013). To control for this, we performed crosstalk correction (Donato et al., 2013). After correction, the number of covered pathways dropped by approximately half (Figure S2A), but eADAGE still covered significantly more pathways than corADAGE (t-test p-value of 0.02) and ADAGE (t-test p-value of 1.29e-05). We subsequently evaluated each method’s coverage of GO biological processes (GO-BP) and found consistent results (Figure S2B). eADAGE integrated multiple models to more broadly capture pathway signals embedded in diverse gene expression compendia.

We next evaluated how specifically and completely signatures learned by the models capture known biology. We use each gene signature’s FDR corrected p-value for enrichment of a KEGG/GO term as a combined measure, as it captures both the sensitivity and specificity. If a pathway was significantly associated with multiple gene signatures in a model, we only considered its most significant association. We found that 71% of KEGG and 79% of GO-BP terms were more significantly enriched (had lower median p-values) in corADAGE models when compared to individual ADAGE models. This increased to 87% for KEGG and 81% for GO-BP terms in eADAGE models. We also directly compared eADAGE and corADAGE by this measure and observed that 74% of KEGG and 61% of GO-BP terms were more significantly enriched in eADAGE. We have found that different pathways were best captured at different model sizes (Figure 2C). We next compared the 300-node eADAGE model to ADAGE models with different number of nodes. Although the 300-node eADAGE models were constructed only from 300-node ADAGE models, we found that 69% of KEGG and 69% of GO-BP terms were more significantly enriched (i.e. lower median p-values) in eADAGE models than ADAGE models of any size, including those with more nodes than the eADAGE models. Three example pathways that are best captured either when model size is small, large, or in the middle are all well captured in the 300-node eADAGE model (Figure 2C). These results demonstrate that eADAGE’s ensemble modeling procedure is effective in capturing consistent signals across models and filtering out noise. Thus, eADAGE more completely and precisely captures the gene expression signatures of biological pathways.

We designed eADAGE to provide a more robust analysis framework than individual ADAGE models. To assess this, we examined the percentage of models that covered each pathway (coverage rate) between ADAGE and eADAGE. The pathways covered by each individual ADAGE model were highly variable. Most KEGG pathways were covered by less than half of individual models but more than half of eADAGE models (Figure 2D), suggesting that eADAGE models were more robust than individual ADAGE models. Subsequent evaluations of GO-BP were consistent with this finding (Figure S2C). We excluded KEGG/GO terms always covered by both individual ADAGE and eADAGE models and observed that 69% of the remaining KEGG and 71% of the remaining GO terms were covered more frequently by eADAGE than ADAGE. This suggests that their associations are stabilized via ensemble construction. In summary, these comparisons of eADAGE and ADAGE reveal that not only are more pathways captured more specifically, but also those that are captured are captured more consistently.

Principal component analysis (PCA) and independent component analysis (ICA) have been previously used to extract biological features and build functional gene sets (Alter et al., 2000; Chen et al., 2008; Engreitz et al., 2010; Frigyesi et al., 2006; Gong et al., 2007; Lutter et al., 2009; Ma and Kosorok, 2009; Raychaudhuri et al., 2000, 2000; Roden et al., 2006). We performed PCA and generated multiple ICA models from the same *P. aeruginosa* expression compendium and evaluated their KEGG/GO term coverage following the same procedures used for eADAGE. eADAGE substantially and significantly outperforms PCA in terms of pathway coverage (Figure 2E). Between eADAGE and ICA, we observed that eADAGE represented KEGG/GO terms more precisely than ICA. Specifically, among terms significantly enriched in either approach, 68%

KEGG and 71% GO terms exhibited more significant enrichment in eADAGE. Increasing the significance threshold for pathway coverage demonstrates the advantage of eADAGE (Figure 3D and Figure S2D).

**Figure 3:**
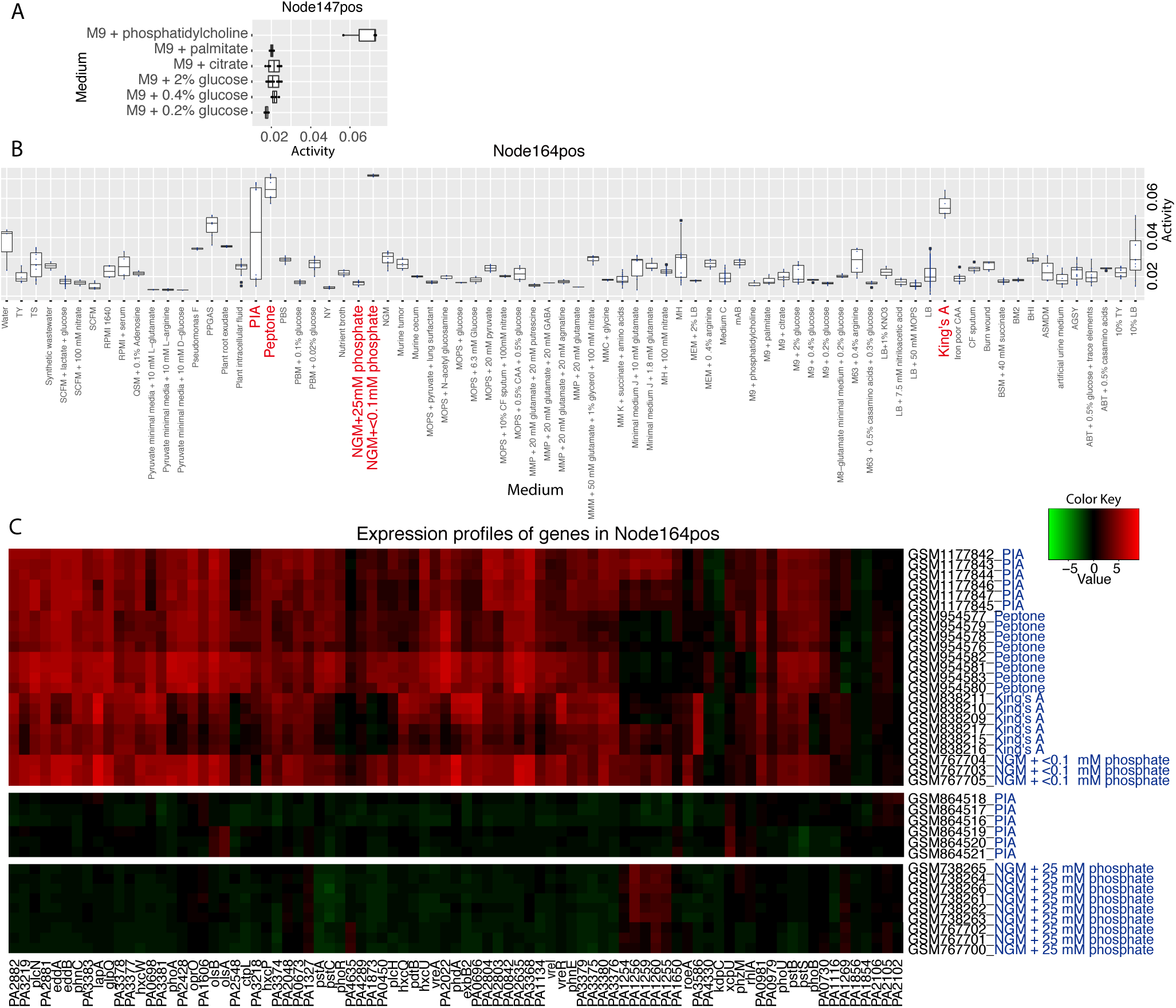
eADAGE signatures that show medium-specific patterns. A Activity of Node147pos in M9-based media. Its activity is high in M9 with phosphatidylcholine but low in other M9-based media. A Activity of Node164pos in all media. NGM+<0.1phosphate, peptone, and King’s A media have evident elevation in Node164pos’s activity. PIA medium show a wide range in Node164pos’s activity. All other media have very low activities. B Expression heatmaps of genes in Node164pos across samples in NGM+<0.1phosphate, peptone, King’s A, and PIA media. Heatmap color range is determined by the Z-scored gene expression of all samples in the compendium. These genes are highly expressed in all samples grown on NGM + <0.1mM phosphate, peptone, King’s A, and half of samples on PIA, but not expressed in samples grown on NGM + 25mM phosphate.

Pathway databases provide a means to compare unsupervised methods for signature discovery. Not all pathways will be regulated at the transcriptional level, but those that are may be extracted from gene expression data. The unsupervised eADAGE method revealed signatures that corresponded to *P. aeruginosa* KEGG/GO terms better than PCA, ICA, ADAGE, and corADAGE. It had higher pathway coverage (breadth), covered pathways more specifically (depth), and more consistently (robustness) than existing methods.

### Elucidating functional signatures that are indicative of growth medium

For biological evaluation, we built a single new eADAGE model with 300 nodes. The model’s weight matrix (Table S2) and all gene signatures (Table S3) are provided. For each signature, we calculated its activity in each sample (see Methods, Table S4). A high activity indicates that the majority of genes in the signature are highly expressed in the sample.

Analysis of differentially expressed genes is widely used to analyze single experiments, but crosscutting signatures are required to reveal general response patterns from large-scale compendia. Signature-based analyses can suggest mechanisms such as crosstalk and novel regulatory networks. However, in order for this to be effective, these signatures must be robust and comprehensive. By capturing biological pathways more completely and robustly, eADAGE enables the analysis of signatures, including those that don’t correspond to any KEGG pathway, across the entire compendium of *P. aeruginosa*.

Gene expression experiments have been used to investigate a diverse set of questions about *P. aeruginosa* biology, and these experiments have used many different media to emphasize different phenotypes. Our manual annotation showed that 78 different base media were used across the gene expression compendium (Table S1). While the compendium contains 125 different experiments, it is exceedingly rare for investigators to use multiple base lab media within the same experiment. There were only two examples in the entire compendium (Table S1). Other than LB, which is used in 43.6% (458/1051) of the samples in the compendium, most media are only represented by a handful of samples.

To provide an illustrative example of cross-experiment analysis, we examined signature activity across the six experiments in a base of M9 minimal medium (Miller, 1972), which used six different carbon sources. Node147pos was highly active in phosphatidylcholine compared to all other media (Figure 3A). This node was significantly enriched for the GO terms choline catabolic process (FDR q-value of 2.9E-11) and glycine betaine catabolic process (FDR q-value of 4.6E-20). Of all signatures, it had the largest overlap with the regulon of GbdR, the choline-responsive transcription factor (Hampel et al., 2014) (FDR q-value of 2.5E-47), suggesting that choline catabolism is active in this medium. Consistent with this, phosphatidylcholine, but not palmitate, citrate, or glucose, serves as a source of choline for *P. aeruginosa* (Wargo et al.,2011, 2009). Importantly, while Node147pos was differentially active within a single experiment containing samples in phosphatidylcholine and palmitate (E-GEOD-7704), it was also identifiable in comparisons to samples grown in M9 medium with different carbon sources in experiments performed in different labs at different times. This illustrates how medium-specific signatures can be identified without experiments designed to explicitly test the effect of a specific medium component on gene expression.

### Distinct aspects of the response to low phosphate are captured among the most active signatures

To broadly examine signatures across all media, we calculated a medium activation score for each signature-medium combination. This score reflected how a signature’s activity in a medium differed from its activity in all other samples (Figure S3, see methods for details). Table S5 lists signatures with activation scores in a specific medium above a stringent threshold. A signature could be active in multiple media (Figure S3), so we averaged their activation scores when this occurred. Table S6 lists signatures that are most active in a group of media (a complete list of signature-media group associations is in Table S7).

The two signatures with the highest pan-media activation scores were Node164pos and Node108neg (Table S6). To evaluate the basis for the high activation scores, we examined their underlying activities across all media (Node164pos is shown Figure 3A), and found that both were highly active in King’s A medium, Peptone medium, and NGM+<0.1mM phosphate (NGMlowP), but not in NGM+25mM phosphate (NGMhighP). The activity differences between NGMlowP and NGMhighP suggested that these signatures respond to phosphate levels. The other two media (Peptone and King’s A) in which Node164pos had high activity also had low phosphate concentrations (0.4 mM) relative to other media. For example, commonly used LB has a phosphate concentration of ∼4.5 mM (Bertani, 2004) and many others have concentrations above 20 mM.

KEGG pathway enrichment analysis of Node164pos genes showed enrichment in phosphate acquisition related pathways (Table S6). One Node164pos gene encodes PhoB, a transcription factor in the PhoR-PhoB two-component system that responds to low environmental phosphate in *P. aeruginosa* (Bielecki et al., 2015; Blus-Kadosh et al., 2013; Santos-Beneit, 2015). Further, Node164pos is the signature most enriched for a previously defined PhoB regulon (FDR q-value of 8.1e-29 in hypergeometric test).

Expression levels of genes in Node164pos are higher in Peptone, King’s A, and NGMlowP than in NGMhighP (Figure 3B), including *phoA* which encodes alkaline phosphatase, an enzyme whose activity can be monitored using a colorimetric assay. As expected, PhoA was activated when phosphate concentrations were low (Figure 4A). Furthermore, PhoA activity was dependent on PhoB and the PhoB-activating histidine kinase PhoR, consistent with published work (Bielecki et al., 2015). Notably, PhoA activity was evident on King’s A and Peptone (Figure 4B). Although King’s A and Peptone are not considered to be phosphate-limited media, these results provide striking evidence that they induced PhoB activity as predicted by Node164pos’s signature-medium relationship.

**Figure 4:**
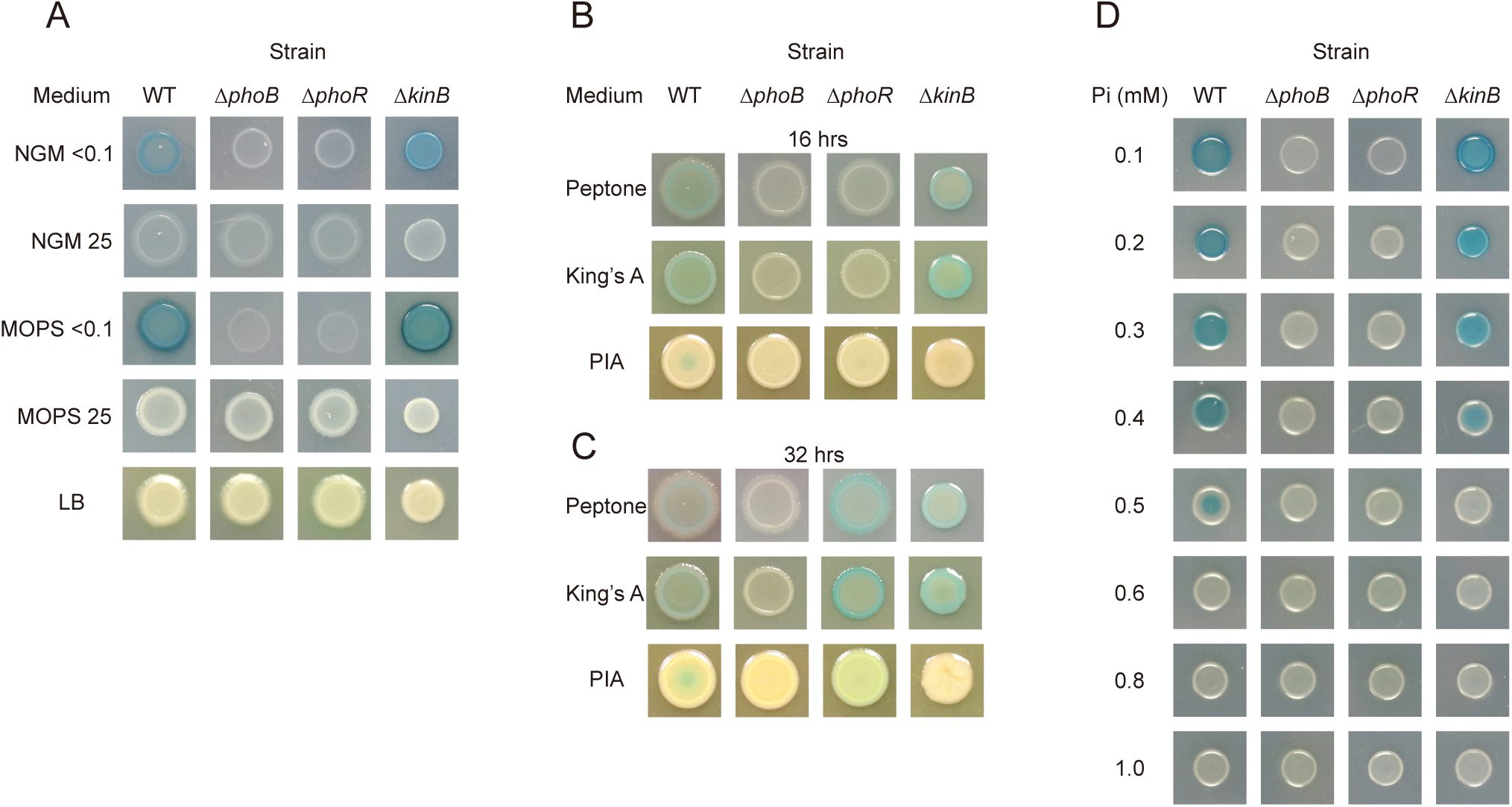
PhoA activity, as seen by the colorimetric BCIP assay in various media. A PhoA activity, as seen by the blue-colored product of BCIP cleavage, is dependent on low phosphate concentrations, phoB, phoR and, in NGM, kinB. B PhoA is active in King’s A, Peptone and PIA and is dependent on phoB and phoR on King’s A and peptone but dependent on kinB as well on PIA at 16 hours. C PhoA is active in King’s A, Peptone and PIA and is dependent on phoB, but no longer phoR, while still dependent on kinB on PIA after 32 hours. D PhoA activity is dependent on phosphate concentrations < 0.6 mM, phoB, phoR and kinB as well at 0.5 mM phosphate in MOPS. Concentration 0.2 mM (not shown) mimics 0.1mM and concentrations 0.7mM - 0.9mM (not shown) mimic 1.0 mM.

While Node108neg is not significantly associated with phosphate acquisition-related KEGG pathways, it is enriched for the PhoB regulon (FDR q-value of 5.2e-9 in hypergeometric test, Table S6) and shares over half of its thirty-two genes with Node164pos. Six of the seven PhoB-regulated genes present in Node108neg are also regulated by TctD, a transcriptional repressor described by Haussler and colleagues (Bielecki et al., 2015). Therefore, Node108neg primarily represents genes that are both PhoB-activated and TctD-repressed. Subsequent analyses found that Node108neg was the most differentially active signature between a *ΔtctD* strain and the wild type in an RNAseq experiment (E-GEOD-64056). Importantly, eADAGE learned this TctD regulon even though the expression compendium did not contain any samples of *tctD* mutants. This demonstrates the utility of eADAGE in learning regulatory programs uncharacterized by KEGG.

We evaluated whether the PhoB and TctD signals were also extracted by PCA, ICA, or ADAGE. ICA and ADAGE captured signatures enriched of the PhoB regulon less than those of eADAGE (Table S8). PCA captured a strong PhoB signal in its 19th principal component. However, it did not learn the subtler TctD signal. In summary, the other methods were able to capture some of this signature but in a manner that was less complete or failed to separate TctD.

### Cross-compendium analysis of Node164pos activity reveals a role for the histidine kinase KinB in the regulation of PhoB

Interestingly, Node164pos activity exhibited a wide spread in PIA medium, with six samples having high activities and the other six having low activities (Figure 3A). All of the strains in which Node164pos was low were from a study that used a PAO1 *kinB*::Gm^R^ mutant background (Damron et al., 2012). The PIA-grown samples with high Node164pos activity used a PAO1 strain with kinB intact (Damron et al., 2013) leading us to propose that KinB may be a regulator of PhoB on PIA. We confirmed that PhoA activity dependents on PhoB, PhoR, KinB on PIA medium (Figure 4B) as illustrated by the fact that a screen of 63 histidine kinase in-frame deletion mutants (Table S9) found only *ΔphoR* and *ΔkinB* had no PhoA activity on PIA, like the *phoB* mutant. These kinases appear to regulate PhoB non-redundantly and to different extents in PIA, as the *ΔphoR* mutant regained PhoA activity at later time points but the *ΔkinB* mutant did not (Figure 4C).

Although the phosphate concentration of PIA (0.8mM) is lower than that of rich media such as LB (∼4.5mM), it is higher than that of Peptone and King’s A (0.4mM). Therefore, we tested whether a moderately low level of phosphate provokes KinB regulation of PhoA. Like in PIA, we found that PhoA activity was evident at concentrations up to 0.5 mM phosphate in MOPS medium in the wild type, but only at lower concentrations in the *ΔkinB* strain suggesting that KinB plays a role at intermediate concentrations (Figure 4D). To our knowledge, KinB has not been previously implicated in the activation of PhoB.

In summary, eADAGE effectively extracted biologically meaningful features, accurately indicated their activity in multiple media spanning numerous independent experiments, and revealed a novel regulatory mechanism. By summarizing gene-based expression information into biologically relevant signatures, eADAGE greatly simplifies analyses that cut across large gene expression compendia.

## Discussion

Our eADAGE algorithm combines multiple ADAGE models into one ensemble model to address model variability due to stochasticity and local minima. The algorithm is inspired by consensus clustering, which reconciles the differences in cluster assignments in multiple runs. Comparable approaches have also been applied for ICA, where researchers have used the centrotypes in clustering multiple models as the final model (Himberg et al., 2004). The ICA centrotype approach for ADAGE corresponds to corADAGE, and our comparison of eADAGE and corADAGE shows that eADAGE not only covers more biological pathways, but also results in cleaner representations of biological pathways. This direct comparison suggests that placing particular emphasis on the genes most associated with a particular feature may be a useful property for other unsupervised feature construction algorithms. While our results demonstrate that this ensemble process can help improve the biological interpretability of neural networks, we do not expect it to increase prediction accuracies in supervised learning problems.

eADAGE revealed patterns that were detectable from a large data compendium containing experiments performed in 78 different media but that were not necessarily evident in individual experiments. For example, one eADAGE signature revealed media in which *P. aeruginosa* had high PhoB activity. PhoB is a global regulator, and understanding its state in different media can provide important insight into medium-specific phenotypes. King’s A and PIA, on which the PhoB signature was active, are known to stimulate robust production of colorful secondary metabolites (King et al., 1954) called phenazines. Separate studies have shown that PhoB can influence phenazine levels (Jensen et al., 2006). Future studies will reveal whether or not the low phosphate levels in these media contribute to this characteristic phenotype. We expect that other signatures extracted from the compendium by eADAGE will serve as the basis for additional work in which the patterns are not only examined but also validated.

We also uncovered a subtle aspect of the phosphate starvation response that depends on KinB, a histidine kinase not previously associated with PhoB. Bacterial two-component systems are often insulated from each other (Podgornaia and Laub, 2013). Though sensor kinase/response regulator cross-talk has been hypothesized as a mechanism of explaining the complexity of signaling networks (Fisher et al., 1995; Ninfa et al., 1988), it is challenging to find conditions where two kinases are needed for full response regulator activation (Verhamme et al., 2002). We propose that moderate levels of phosphate, like those in PIA, provide a niche for crosstalk: the activity of PhoR is low enough that the interaction with KinB is needed for full PhoB activity on this medium. Together, PhoR and KinB may enable a more sensitive and effective response to phosphate limitation. Alternatively, KinB may influence PhoB activity indirectly by regulating activities that affect PhoB levels, phosphorylation state, or protein-protein interactions. This relationship was not observed in experiments designed to perturb this process, which use high and very low phosphate concentrations. Instead, eADAGE analysis of Pseudomonas aeruginosa transcriptomic measurements across multiple experiments in different media were required to reveal this nuanced mechanism.

Existing public gene expression data compendia for more than one hundred organisms are of sufficient size to support eADAGE models (Greene et al., 2016). Cross-compendium analyses provide the opportunity to efficiently use existing data to identify regulatory patterns that are evident across multiple experiments, datasets, and labs. To tap this potential, we will require algorithms like eADAGE that robustly integrate these diverse datasets in a manner that is not tied to only aspects of biology that are well understood. Furthermore, while public compendia tend to be dominated by expression data, autoencoders have also been successfully applied to datasets based on large collections of electronic health record (Beaulieu-Jones et al., 2016; Miotto et al., 2016). Within the health records space, these methods are particularly effective at dealing with missing data (Beaulieu-Jones et al., 2016; Beaulieu-Jones and Moore, 2017). These features, along with their unsupervised nature, make DAs a promising approach for the integration of heterogeneous data types. We find that ensembles of DAs construct clearer features that more robustly capture biological processes. Ultimately, we expect unsupervised algorithms to be most helpful when they lead users to discover new underlying mechanisms, which require models that are accurate, robust, and interpretable.

## Acknowledgements

This work was supported in part by a grant from the Gordon and Betty Moore Foundation (GBMF 4552) to CSG. This work was supported by National Institutes of Health (NIH) grant RO1-AI091702 to DAH. MTL is an investigator of the Howard Hughes Medical Institute. This work was supported by a pilot grant from the Cystic Fibrosis Foundation (STANTO15R0) to CSG and DAH. The authors would like to thank Gregory Way and René Zelaya for helpful code review. The authors also would like to thank Anastasia Baryshnikova for providing critical feedback on a preprint of this work.

## Author contributions

JT, DAH and CSG conceived and designed the research. JT, GD and KMC performed computational analyses. GD, KAL and CEP performed molecular experiments. KC, BP and MTL constructed and contributed the histidine kinase knock out collection. JT, GD, KMC, DAH and CSG wrote the manuscript, and KAL, CEP, KMC, KD, BP and MTL provided critical feedback.

## METHODS

### Data processing

We followed the same procedures for data collection, processing, and normalization as (Tan et al., 2016b) and updated the *P. aeruginosa* gene expression compendium to include all datasets on GPL84 platform from the ArrayExpress database (Rustici et al., 2013) as of 31 July 2015. This *P. aeruginosa* compendium contains 125 datasets with 1051 individual genome-wide assays. Processed expression values of the ΔtctD RNAseq dataset were downloaded from ArrayExpress (E-GEOD-64056) and normalized to the range of the compendium using TDM (Thompson et al., 2016). We provide the *P. aeruginosa* expression compendium (Dataset S1) along with all the code used in this paper (Tan et al., 2016a). The eADAGE repository is also tracked under version control at https://bitbucket.org/greenelab/eadage.

### Construction of ADAGE models

We constructed ADAGE models as described in (Tan et al., 2016b). To summarize the process and outputs, we constructed a denoising autoencoder for the gene expression compendium. Denoising autoencoders model the data in a lower dimension than the input space, and the models are trained with random gene expression measurements set to zero. Thus an ADAGE model must learn gene-gene dependencies to fill in this missing information. Once the ADAGE model is trained, each node in the hidden layer contains a weight vector. These positive and negative weights represent the strength of each gene’s connection to that node.

## Gene signatures as sign-specific high-weight gene sets

In previous work (Tan et al., 2016b) we defined high-weight (HW) genes as those in the extremes of the weight distribution on the positive or negative side of a node. Here, we use a more granular definition that accounts for sign specificity. Each node’s gene weights are approximately normal and centered at zero in ADAGE models (Tan et al., 2016b, 2015). We defined positive HW genes as those that were more than 2.5 standard deviations from the mean on the positive side, and negative HW genes as those that were more than 2.5 standard deviations from the mean on the negative side. After this split, a model with n nodes provides 2n gene signatures. Because a node is simply named by the order that it occurs in a model, we named two gene signatures derived from one node as “NodeXXpos” and “NodeXXneg”.

## KEGG pathway and GO-BP term enrichment analysis

To evaluate the biological relevance of gene signatures extracted by an ADAGE model, we tested how they related to known KEGG pathways (Kanehisa and Goto, 2000). We tested a signature’s association with each KEGG pathway using hypergeometric test and corrected the p-value by the number of KEGG pathways we tested following the Benjamini-Hochberg procedure. We used a false discovery rate of 0.05 as the significance cutoff. The same procedure was repeated using GO-BP terms. We downloaded biological process GO terms from pseudomonas.com and only used manually curated terms. For KEGG and GO terms, we only considered terms with more than 5 genes and less than 100 genes as meaningful pathways or processes.

Genes can be annotated to multiple pathways. To control for this effect in our analysis, we also performed a parallel analysis after applying crosstalk correction as described in (Donato et al., 2013). This approach uses expectation maximization to map each gene to the pathway in which it has the greatest predicted impact. A gene-to-pathway membership matrix, defined using KEGG pathway annotations, initially makes the assumption that each gene’s role in all of its assigned pathways remains constant independent of context. We then applied pathway crosstalk correction using genes’ weights for each node in the ADAGE model. We used the expectation maximization algorithm to maximize the log-likelihood of observing the membership matrix given each node’s weight vector. This process inferred an underlying gene-to-pathway impact matrix and iteratively estimated the probability that a particular gene g contributed the greatest fraction of its impact to some pathway P. Upon convergence, we assigned each gene to the pathway in which it had the maximum impact. The resulting pathway definitions do not share genes. We then used these corrected definitions for an analysis parallel to the KEGG process described above.

## Reconstruction error calculation

The training objective of ADAGE is to, given a sample with added noise, return the originally measured expression values. The error between the reconstructed data and the initial data is the ‘reconstruction error.’ To summarize the difference over all genes we used cross-entropy between the original sample and the reconstruction, which has been widely used with these methods and in this domain (Tan et al., 2016b; Vincent et al., 2008). This matches the statistic used during training of the model. To calculate reconstruction error for a model, we use the mean reconstruction error across samples.

## Model size and sample size heuristics

One important parameter of a denoising autoencoder model is the number of nodes in the hidden layer, which we refer to as the model size. To evaluate the impact of model size and choose the most appropriate size, we built 100 ADAGE models at each model size of 10, 50, 100, 200, 300, 500, 750, and 1000, using different random seeds. The random seed determines the initialization values in the weight matrix and bias vectors in ADAGE construction, so different random seeds will result in models that reach different local minima. Other training parameters were set to the values previously identified as suitable for a gene expression compendium (Tan et al., 2015). In total, 800 ADAGE models, i.e. 100 at each model size, were generated in the model size evaluation experiment.

To evaluate the impact of sample size on the performance of ADAGE models, we randomly generated subsets of the *P. aeruginosa* expression compendium with sample size of 100, 200, 500, and 800. We then trained 100 ADAGE models at each sample size, each with a different combination of 10 different random subsets and 10 different random training initializations. To evaluate each model, we randomly selected 200 samples not used during training as its testing set. We performed this subsampling analysis at model size 50 and 300. In total, 800 ADAGE models were built in the sample size evaluation experiment.

The impacts of model size and sample size on model selection were evaluated in the supplement (Figure S4). For subsequent steps, we set the model size to 300 because it was the size that was best supported in the current *P. aeruginosa* compendium by this evaluation.

## Construction of eADAGE models

We constructed ensemble ADAGE (eADAGE) models by combining many individual ADAGE models in to a single model. For each eADAGE model we combined 100 individual ADAGE models. The 100 models were trained with identical parameters but distinct random seeds. For an eADAGE model of size 300, we trained 100 individual models with 300 nodes each, which provided 30000 total nodes. Each node has a weight vector. We have previously observed that high-weight genes provided the most information to each node (Tan et al., 2016b), so we calculated a weighted Pearson correlation between each node’s weight vectors. Our weighted Pearson correlation used (| node1 weight| + | node2 weight|)/2 as the weight function for each gene. We compared this to an unweighted Pearson correlation (corADAGE) as well a baseline ADAGE model.

After calculating correlation (weighted for eADAGE and unweighted for corADAGE), we converted the correlation to distance by calculating (1-correlation)/2. This provided a 30000*30000 distance matrix storing distances between every two nodes. We clustered this distance matrix using the Partition Around Medoids (PAM) clustering algorithm (Park and Jun, 2009).We implemented clustering in R using the ConsensusClusterPlus package (Wilkerson and Hayes, 2010) from Bioconductor with the ppam function from Sprint package to perform parallel PAM (Piotrowski et al., 2011). We set the number of clusters to match the individual ADAGE model (e.g. 300) allowing for direct comparison between the eADAGE and ADAGE methods.

Clustering assigned each node to a cluster ranging from 1 to 300. We combined nodes assigned to the same cluster by calculating the average of their weight vectors. These 300 averaged vectors formed the weight matrix of the eADAGE model. Because the ensemble model is built from the weight matrices of individual models, it does not have the parameters that form the bias vectors. We built 10 eADAGE and 10 corADAGE models from 1000 ADAGE models with each ensemble model built upon 100 different individual models. The individual eADAGE model used for biological analysis in this work was constructed with random seed 123, which was arbitrarily chosen before model construction and evaluation.

## PCA and ICA model construction

We constructed PCA and ICA models and defined each model’s weight matrix following the same procedures in (Tan et al., 2016b). To compare with the 300-node eADAGE, we generated models of matching size (300 components). For ICA, we evaluated 10 replicates. PCA provides a single model. PCA and ICA models were evaluated through the KEGG pathway enrichment analysis described above.

## Activity calculation for a gene signature

We calculated a signature’s activity for a specific sample as *A* = *W* · *E/N*, in which W is a vector of genes’ absolute weights in that signature, E is a vector of genes’ expression values after zero-one normalization in that sample, and N is the number of genes. It can be viewed as an averaged weighted sum of genes' expression levels for genes in the signature. We normalized a signature’s activity by the number of genes (N) in that signature, because different signatures have different number of genes. We use gene’s absolute weight value in activity calculation to keep activity positive. In this way, a high activity indicates that majority of genes in the signature are highly expressed in the sample and a low activity indicates that majority of genes in the signature are lowly expressed in the sample.

## Media annotation of the ****P. aeruginosa**** compendium

A team of *P. aeruginosa* biologists annotated the media for all samples in the compendium by referring to information associated with each sample in the ArrayExpress (Rustici et al., 2013) and/or GEO (Edgar, 2002) databases and along with the original publication, if reported. Each sample was annotated by two curators separately. Conflicting annotations, if they occurred, were resolved by a third curator. The media annotation for all samples in the compendium were provided in Table S1.

## Identification of signatures activated across media

We calculated an activation score to identify gene signatures with dramatically elevated or reduced activity in a specific medium. We grouped samples by their medium annotation. For each gene signature and medium combination, we calculated the absolute difference between the mean activity of the signature for samples in that medium as well as the mean activity across the remainder of samples in the compendium. We divided this difference in the means by the range of activity for all samples across the compendium. This score captures the proportion by which the mean activity in a medium differs relative to the total difference across the compendium. We termed this ratio the activation score.

To identify the most specifically active signatures for each medium, we constructed a table for all pairs with an activation score greater than or equal to 0.4 (Table S5). This was highly stringent: it captured only the top 2.4% of the potential signature-medium pairs (Figure S9). To identify pan-media signatures, we limited signatures to those that were active in multiple media (greater or equal to 0.4) and averaged their activation scores (Table S7). These signatures exhibit parallel patterns for multiple media across multiple distinct experiments.

## Definition of the PhoB regulon

A PhoB regulon for the PAO1 genome was adapted from the PhoB regulon of PA14 in (Bielecki et al., 2015) in order to be comparable to models built with PAO1 genome. Of the 187 genes in the PA14 regulon, 160 were in the PAO1 reference genome (www.pseudomonas.com).

## Strains and Media

Strains used were WT, ΔphoB (DH2633, O’Toole lab collection), ΔphoR (DH2516) and ΔkinB (DH2517), all in the PA14 background. All strains were maintained on LB with 1.5% agar and grown at 37 °C. For cross-media and phosphate concentration comparisons, BCIP assays (see methods below) were conducted on different base media with 1.5% agar (Fisher): King’s A (Pancreatic Digest of Gelatin (Difco) 20g/L; MgCl2 1.4g/L; K2SO4 10g/L; Glycerol 10ml/L) (King et al, 1954), LB (Tryptone (Fisher) 10g/L; Yeast Extract (Fisher) 5g/L; NaCl 5g/L) (Bertani, 2004), MOPS (morpholinepropanesulfonic acid 40mM; Glucose 20 ml/L; K2SO4 2.67mM; K2HPO2 0mM, 25mM or 0.1 - 1 mM) (Neidhardt et al., 1974), NGM (Pancreatic Digest of Gelatin 2.5g/L;

Cholesterol 5mg/L; NaCl 3g/L; MgSO4 1mM; CaCl2 1mM; KCl 25mM; Potassium Phosphate buffer pH6 0 or 25 mM) (Zaborin et al., 2009), Peptone (Pancreatic Digest of Gelatin 10g/L; MgSO4 1.5g/L; K2SO4 10g/L) (Lundgren et al., 2013), Pseudomonas Isolation Agar (PIA, prepared as per instructions, BioWorld).

## BCIP assay

Various media were supplemented with 5-bromo-4-chloro-3-indolyl phosphate (BCIP) DMF solution to a final concentration of 60 μg/mL. BCIP assay plates were inoculated with 5 μl of overnight *P. aeruginosa* culture in LB broth. Colonies were grown for 16 hours at 37 °C then matured at room temperature until imaging. Images were collected 16 and 32 hours post inoculation.

## Screen of a histidine kinase mutant collection

Molecular techniques to construct the histidine kinase (HK) knock out collection were carried out as previously described (Ha et al., 2014). For each strain in the HK collection, a BCIP assay was performed on PIA. Plates were struck with an overnight *P. aeruginosa* culture concentrated two-fold by centrifugation. Plates were incubated at 37 °C 12-16 hours and matured at room temperature for an additional 12-16 hours alkaline phosphatase activity was determined qualitatively, based on blue color.

